# Human single-cell atlas analysis reveals heterogeneous endothelial signaling

**DOI:** 10.1101/2025.08.15.670370

**Authors:** Zimo Zhu, Rongbin Zheng, Yang Yu, Lili Zhang, Kaifu Chen

## Abstract

Endothelial cells (ECs) play complex roles across tissues and vessel types. Yet, systematic investigations of EC heterogeneity in the combined context of vessel type and tissue microenvironment are still largely lacking. We integrated over three million single cells of scRNA-seq datasets in 15 human tissues and found that ECs in some tissues (e.g., heart and kidney) exhibited greater tissue specificity, while others displayed more substantial vessel specificity. We developed a computational pipeline to analyze cell-cell communications (CCC) mediated by metabolites or proteins to explore microenvironmental regulation. Interestingly, our results showed that CCC events involving ECs varied vastly across tissues, highlighting tissue-specific EC interactions. Using topic modeling, we identified CCC patterns, termed CCC topics, representing metabolite- and protein-mediated interactions between ECs and other tissue-resident cells. Most CCC topics exhibited high tissue specificity, potentially explaining microenvironmental regulations for EC heterogeneity. The work systematically investigates EC heterogeneity and provides insights into how EC heterogeneity was regulated across diverse tissue microenvironments.

## Introduction

Endothelial cells (EC) line the cardiovascular vessels throughout the human body, performing essential roles in various biological processes, including tissue homeostasis, immune responses, and vascular health[1–4]. Despite ECs’ common presence, they exhibit remarkable diversity in structure and function that varies by vessel types, anatomic location, and developmental stage [5], demonstrating heterogeneity within the vascular system. Major vessel types include the arterial, venous, capillary, and lymphatic vessels. For example, arterial ECs express markers such as ephrin-B2, which marks arterial identity in the arteries [6]. Venous ECs express EphB4, establishing venous characteristics in the veins [7]. EC also shows differences between tissue types. These differences allow ECs to perform tissue-specific molecular mechanisms, such as cardiac vascular morphogenesis in the heart, gas exchange in the lung, or filtration in the kidney [8]. Studies using single-cell RNA sequencing (scRNA-seq) have shown that ECs exhibit cellular heterogeneity and metabolic plasticity across tissues in mice [9–11] and humans [12–14]. However, there is still a lack of systematic comparison of EC heterogeneity across combined contexts of different vessels and tissues.

It has been reported that the heterogeneity of ECs is regulated by various molecular factors, including transcription factors (TFs) and other signaling molecules [15]. Early in embryogenesis, transcriptional programs driven by TFs, such as GATA2 and ETV2 [16,17], establish the endothelial identity and guide early EC lineage commitment. The differentiated ECs are later refined based on the tissue context in embryonic development. Besides transcriptional regulation, the microenvironment regulates cellular heterogeneity and functions [18]. It is known that microenvironmental factors, including extracellular matrix composition, mechanical forces, and neighboring cell types, modulate gene expression regulation in ECs [19,20]. These regulations make them adapt to the functional needs of respective tissues [20,21]. For instance, ECs in the heart undergo unique shear stress and mechanical forces that upregulate KLF2, promoting an anti-inflammatory gene expression profile [22]. These studies together suggested that the tissue microenvironment is pivotal in establishing tissue-specific EC transcriptomes. Yet, the comprehensive investigation of microenvironmental mechanisms for human tissue-specific ECs remains elusive.

We hypothesized that tissue-specific microenvironments mediated by CCC may regulate human EC heterogeneity across tissue types. CCC is a fundamental process through which cells cross-talk via signaling molecules, such as cytokines and metabolites. The intact CCC network coordinates cell functions, including differentiation, development, and metabolism [23]. For instance, the blood-brain barrier in the brain explicitly influences EC response to neuronal signaling and inflammatory cues [24]. A unique Car4-high EC population was identified in the lung primed to communicate with other alveolar cells during alveolar regeneration after lung injury [25].

This study integrated over 3 million single cells from human scRNA-seq datasets in 15 tissues. First, we systematically assessed the EC heterogeneity across tissues and vessel types. Intriguingly, a higher heterogeneity was observed across vessel types compared to tissue types in most tissues, except the heart, kidney, and intestine. Further, we identified tissue-specific EC topics for metabolite and protein-mediated CCC using a machine learning algorithm, Latent Dirichlet Allocation (LDA). Diving into specific CCC topics, we found that most identified metabolite-sensor and ligand-receptor pairs were tissue-specific and have been reported as vital in carrying out tissue-specific functions. Our work showed a systematic EC heterogeneity analysis across vessel and tissue types. Moreover, the identified tissue-specific CCC topics hold the potential to explain the EC heterogeneity regulated by tissue microenvironmental factors and further provide insights into therapeutic target discovery for endothelial-related diseases.

## Results

### Heterogeneity of EC across tissue and vessel types

We integrated scRNA-seq data from 15 human tissue types (**Supplementary Table 1**) and visualized the EC population using UMAP to investigate the human EC heterogeneity. During the integration, we observed that batch effects may confound the cell clustering as different projects dominate the separation into different clusters (**Supplementary** Fig.1). Exemplified by cells from the heart tissue, the cells in the same tissue type or vessel type from different projects were separated into different clusters (**Fig. 1A, B**). However, the ECs of the same tissue type from different projects were no longer clustered separately after correcting the batch effects (**Supplementary** Fig.2). Instead, ECs became clustered by tissue or EC vessel types (**Fig. 1C, D**). Thereafter, we performed all follow-up analyses based on batch-corrected data. Interestingly, the principal component analysis indicated that all ECs in the heart and kidney, regardless of vessel types, appeared to be very different from those in the other tissue types (**Fig. 1E**). However, most ECs tended to be grouped by vessel types rather than by tissue types, such as the ones grouped into arterial and lymphatic ECs (**Fig. 1E**). To further confirm this observation, we next quantified the EC similarity among tissue and vessel types using unsupervised hierarchical clustering method. Two primary clusters (cluster 1 and cluster 2) were observed, as shown by the largest cophenetic distance based on the height of the dendrogram (**Fig. 1F**). In cluster 1, the distance between ECs was significantly greater within tissue type than within vessel type, indicating that ECs were more similar within vessel type than within tissue type. However, the ECs in cluster 2 showed a larger distance within a vessel type than within a tissue type, meaning that ECs within a tissue type are more similar. These results suggested that the human EC posed significant heterogeneity across vessel and tissue types. Furthermore, some ECs were more heterogeneous across vessel types than tissue types, while some ECs were more heterogeneous across vessel types.

**Figure 1:**
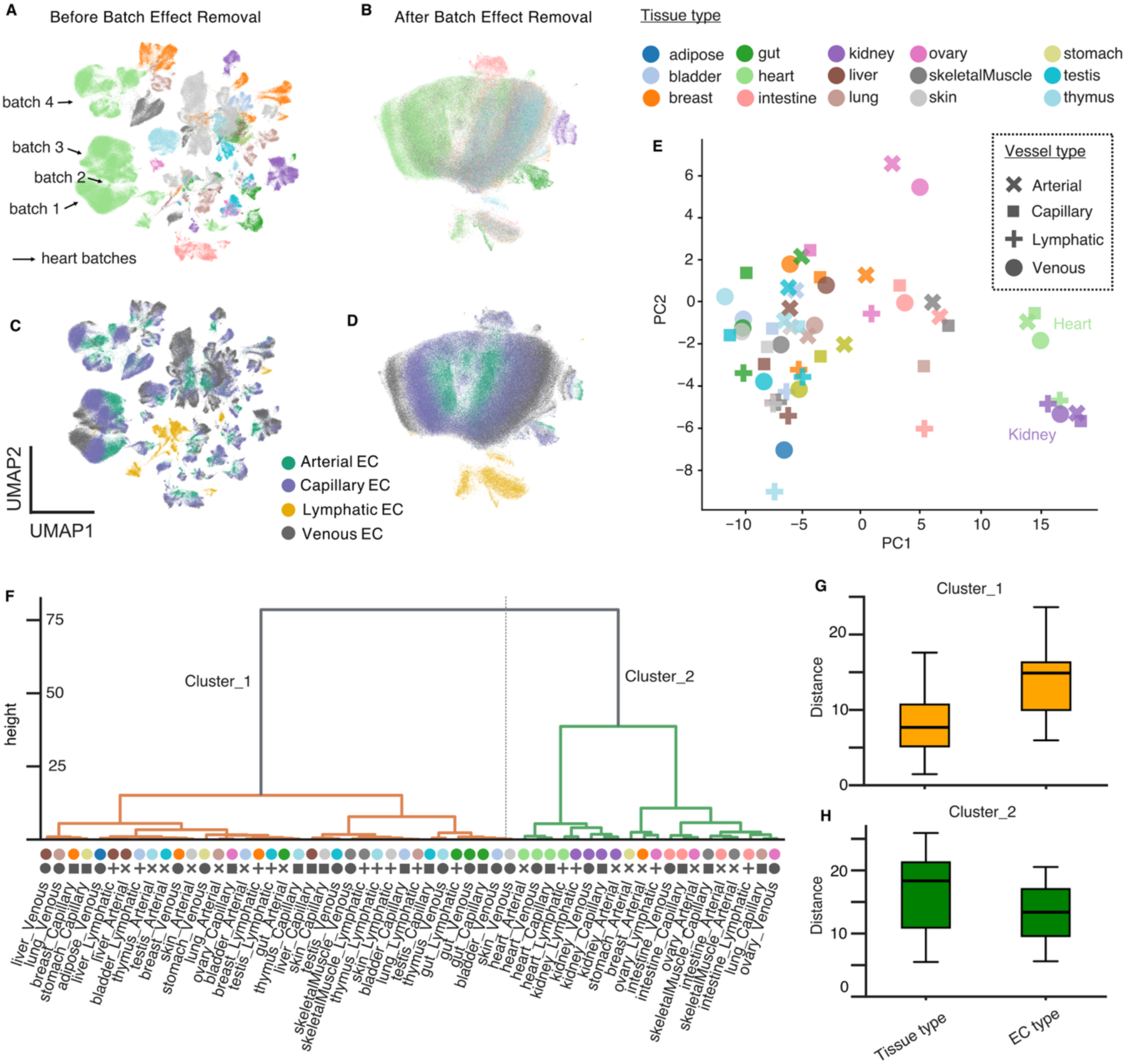
EC demonstrated heterogeneity across tissue and vessel types. (**A-D**) UMAP clustering visualization of EC across tissue and vessel types before (**A**, **B**) and after (**C, D**) batch effect removal, with color-coded for tissue type (**A, C**) and vessel types (**B, D**). (E) PCA projection of EC across tissue type and vessel type. Color-coded for tissue type (consistent with **1A** and **1B**) and shape-indicated EC vessel type. (F) Hierarchical clustering showing tissue type and vessel type on the x-axis and cophenetic distance on the y-axis. The dotted line separated the two major clusters. Colors and shapes were used according to the PCA project in **E**. (**G-H**) Pairwise distances were calculated for cluster 1 on the top (**G)** and cluster 2 on the bottom (**H**). The box on the left shows the results grouping by tissue type and calculating the vessel type distances, and the box on the right shows the results grouping by vessel types and calculating the tissue distances. The median was shown with the bold line in the middle.

### A computational workflow to unravel tissue-specific microenvironmental signaling

While the EC population in cluster 1 was predominantly heterogeneous across vessel types, they still showed substantial heterogeneity across tissue types within a vessel type. This raised the question of how tissue-specific signals were associated with EC tissue heterogeneity. Previous reports show that CCC can regulate tissue-specific EC functions [26]. To study tissue-specific CCC patterns, we developed a computational workflow by integratively analyzing two types of CCC events, namely, metabolite-mediated and protein-mediated CCC (**Fig.2A**). First, cutting-edge bioinformatics tools were implemented to detect CCC events in scRNA-seq data of each tissue. Specifically, MEBOCOST [27] was used for metabolite-mediated CCC detection, while LIANA+ [28,29] was used to detect protein-mediated CCC events. Next, we applied topic modeling, an unsupervised machine learning algorithm, to detect CCC topics among all CCC events across tissues. Topic modeling was extensively used in natural language processing tasks, such as summarizing documents based on groups of words. Here, we defined “document” as tissue and “words” as a collection of CCC events. The output “topics” represent CCC patterns that could be either housekeeping or tissue-specific across tissue types (**Fig. 2B**). The model also outputs two matrices reflecting the probability of a CCC belonging to a topic and the probability of a tissue including a topic, respectively (**Fig. 2C**). We identify tissue-specific CCC topics based on the CCC topic probability vector across tissues. Using probability vectors across CCC events, we further determined if any groups of CCC were highly concentrated on a tissue-specific topic. Finally, the workflow further evaluates gene expression specificities of CCC-related genes across tissue types.

**Figure 2:**
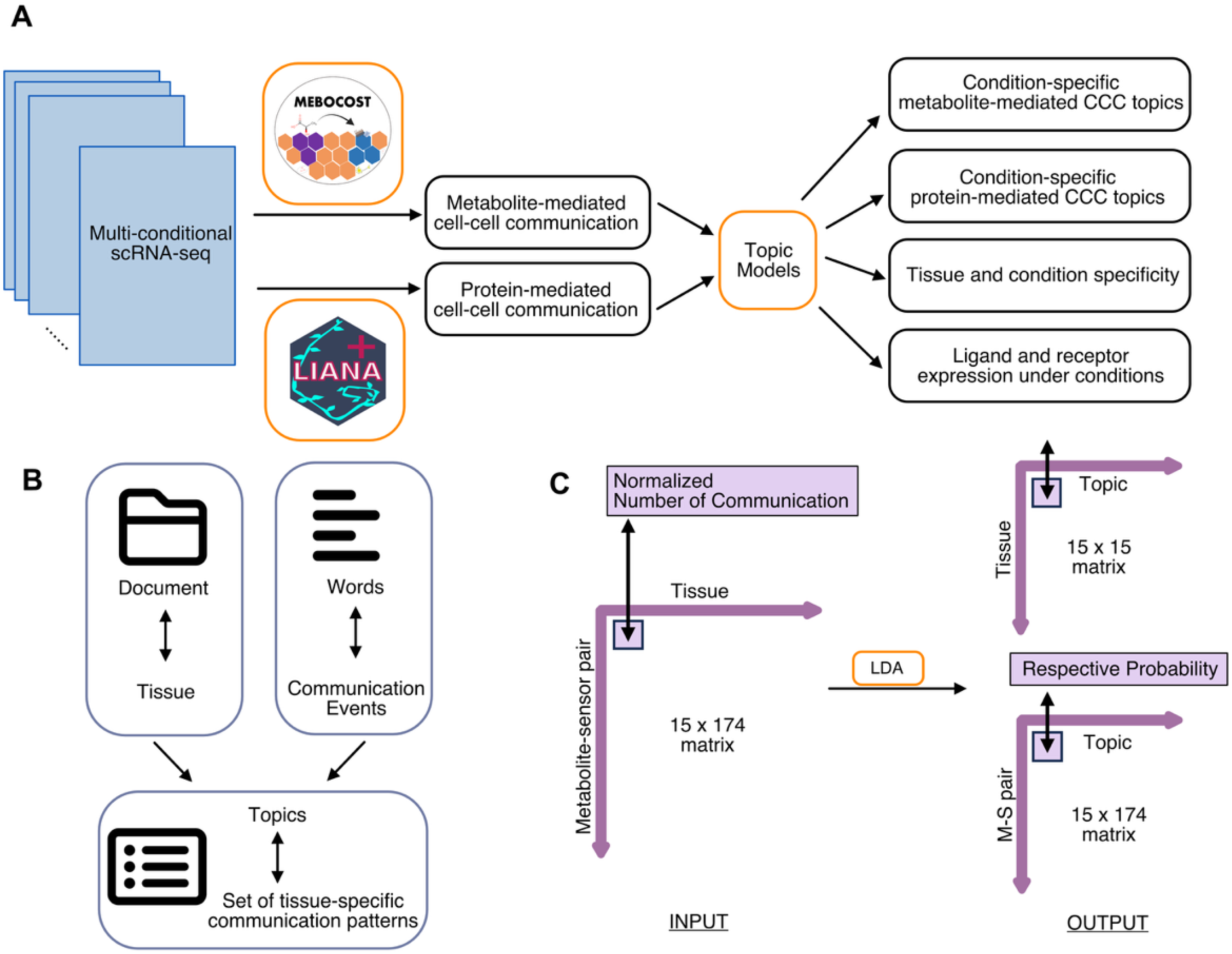
Workflow for processing and analyzing multi-conditional scRNA-seq data. (A) Scheme of analysis design. (B) Explanation of topic modeling under the context of this study. (C) Input and output matrix visualizations for the LDA algorithm.

### Great heterogeneity of EC-related CCC across tissue types

We sought to explore the number of EC-associated CCC events across tissue types. Interestingly, the number of CCC for EC varied across different tissues (**Fig. 3A**). Some tissues, such as the ovary and intestine, showed a much higher number of CCC events than other tissues, such as kidney and adipose. The observation was highly consistent between metabolite-mediated and protein-mediated CCC, suggesting tissue specificity of EC communication with resident cells. Since the number of CCC might be affected by EC population size and tissue cell type number, we computed the EC percentage and total cell type number across different tissues (**Fig. 3B**). Surprisingly, the EC percentage in a tissue showed little correlation (R=-0.05) with the total CCC number (**Fig. 3C**), while the number of cell types was strongly correlated with the number of CCC events in the tissue (R=0.76, **Fig. 3D**). This was consistent with the prior knowledge that the diversity of cell types and their spatial organization within a tissue contribute to the richness of CCC networks [23]. However, the observation of varying CCC numbers across tissue types remained even when it was normalized by the number of cell types in the tissue (**Fig. 3E**). Taken together, these results showed the diverse capability of EC to receive CCC signaling in different tissues, indicating tissue-specific CCC patterns.

**Figure 3:**
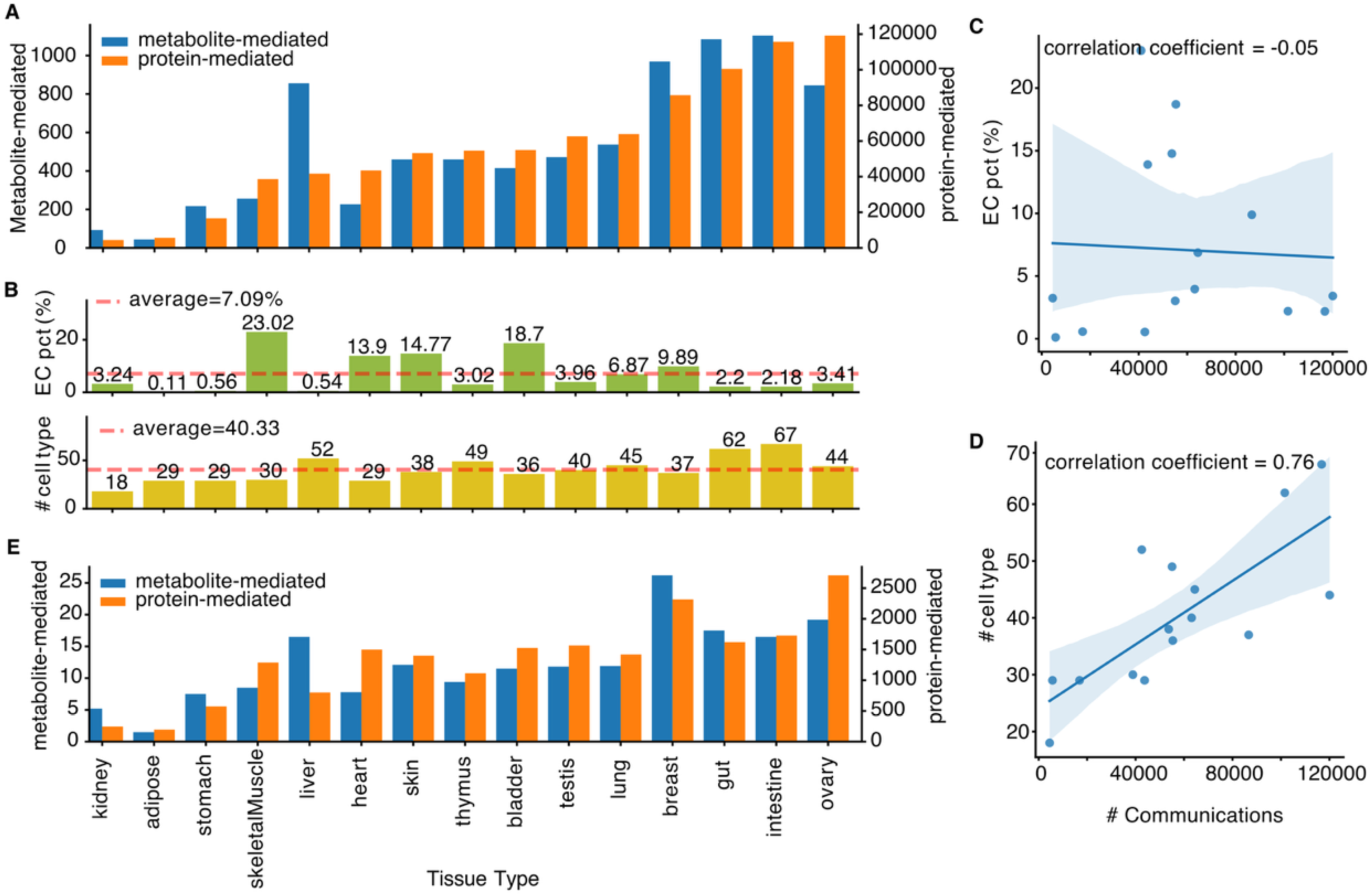
Analysis of metabolite-sensor and ligand-receptor CCC. (A) A bar chart shows the number of communications inferred *in silico*. The blue color indicates the number of metabolite-mediated CCC (refers to the left y-axis scale). The orange color indicates the number of ligand-receptor CCC (refers to the right y-axis scale). (B) Bar chart showing the percentage of EC in the whole cell population and the number of cell types included in a tissue. The red-dotted line indicates the average value under each category. (C) The scatter plot shows the number of communications against EC percentage. Each dot represents a tissue type. The regression line is shown in blue. (D) The scatter plot shows the number of communications against the number of cell types. Each dot represents a tissue type. The regression line is shown in blue. (E) The bar chart shows the number of CCC events of every hundred cell types in each tissue to normalize the effect of total cell type numbers across tissues.

### Topic modeling revealed tissue-specific metabolite-mediated CCC patterns

Functional metabolite-sensing machinery facilitates the crosstalk between the cellular biological network and its tissue microenvironment cues [30]. Recent studies have highlighted dynamic mechanisms with metabolites acting as signaling mediators to modulate processes like inflammation and cancer progression [31]. To study EC tissue-specific heterogeneity under the context of metabolite-mediated CCC, we applied topic modeling using communications detected in each tissue and identified 15 metabolite CCC topics (**Supplementary Table 2**). Of 15 topics, 9 topics were highly tissue-specific. For instance, topic 11 was highly correlated with thymus.

Topic 5 was liver-dominant based on the topic-tissue probability vectors (**Fig. 4A**). Other less tissue-specific topics were also identified, such as topic 7, topic 2, and topic 10, suggesting some shared CCC networks for ECs across different tissues. Investigating each metabolite-sensor pair in each topic, we observed that they were highly correlated with the specificity of CCC topics (**Fig.4B**). To confirm this specificity further, we checked the number of communications inferred by MEBOCOST for each tissue and found a matched pattern. The metabolite-sensor pairs that appeared more frequently in a topic also tend to show a higher number of CCC in the tissue related to that topic (**Fig.4C**). For example, we identified topic 13 to be highly associated with the ovary (**Fig.4A**). Five metabolite-sensor pairs, including the Cholesterol (metabolite)∼RORA (sensor), L-Lysine∼SLC7A1, Ornithine∼SLC7A1, Adenine∼SLC35F5, and L-Cysteine∼SLC43A2, exclusively showed high probability in topic 13 (**Fig.4B**), and showed more CCC events in the ovary (**Fig. 4C**). To inspect the results at the expression level, we further assessed the communication score, which was inferred by the metabolite enzyme and sensor expression. We found that all these pairs were highly expressed in the ovary (**Fig.4D**). Interestingly, L-Lysine∼SLC7A1 and Ornithine∼SLC7A1 were the two pairs that showed the highest specificity as they nearly solely expressed in the ovary but not in other tissue. Consistently, many previous studies have demonstrated the importance of the selected metabolite and sensor in carrying out tissue-specific functions. For example, a study found lysine succinylation (Ksuc) as a newly discovered post-translational modification (PTM) that plays an essential role in ovarian aging [32]. The paper pointed out that in the ovaries of aged mice and those with premature ovarian insufficiency (POI), increased levels of Ksuc were observed. Further, SLC7A1 was reported to be a key player in the ovarian tumor microenvironment, influencing both cancer cell behavior and stromal fibroblast activation [33].

**Figure 4:**
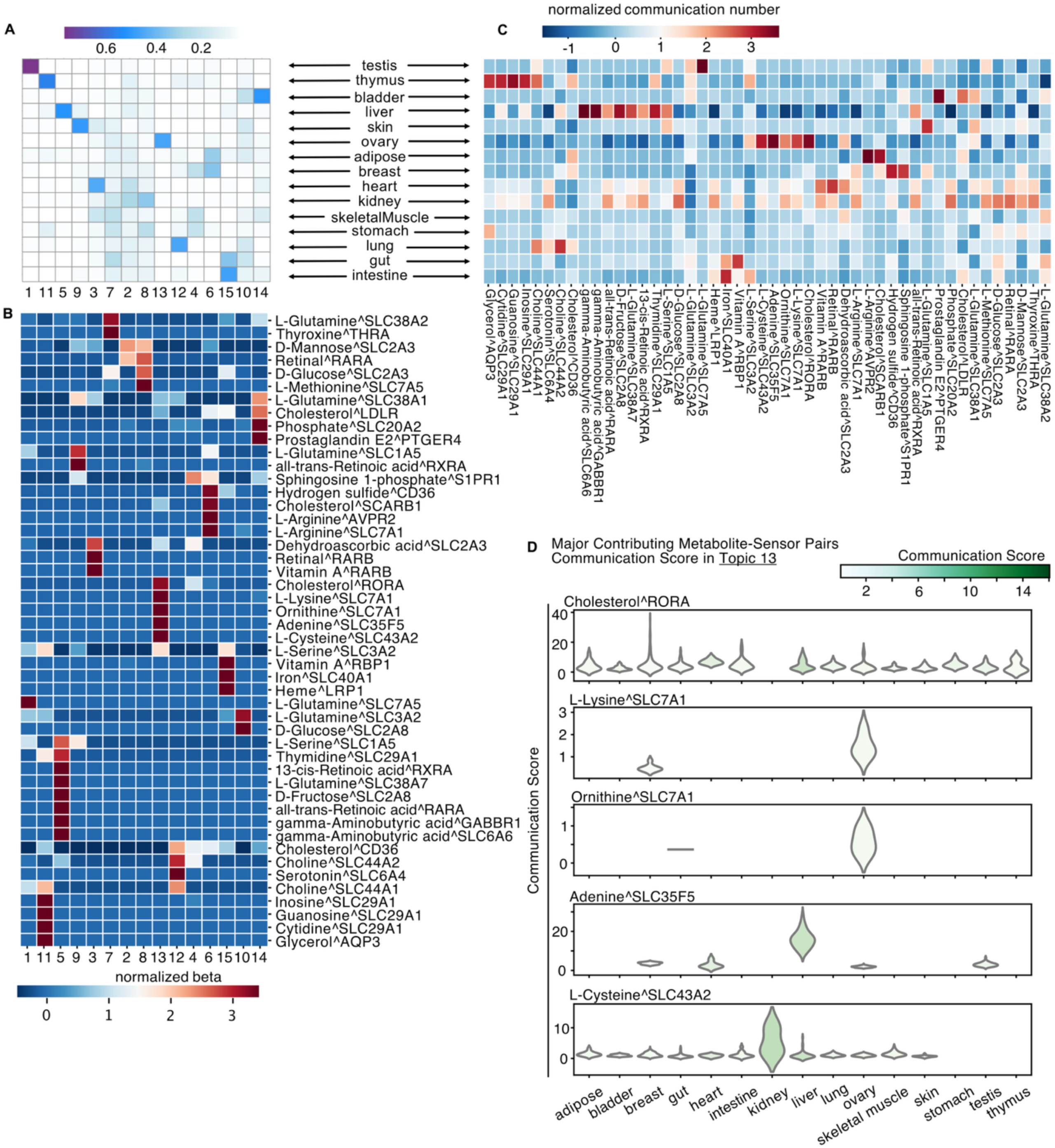
LDA inferred topics of metabolite-mediated CCC networks. (A) Clustered heatmap showing the correlation of the LDA results using MEBOCOST inferred CCC results. The X-axis indicated inferred topics and the y-axis labeled tissue type. The darker the color, the stronger the correlation between the corresponding topic and tissue. (B) Clustered heatmap of the normalized beta score of selected metabolite-sensor pairs (y-axis) in each topic (x-axis). The x-axis label was aligned with that in **4A**. The warmer the color, the higher the specificity, and vice versa. (C) Clustered heatmap of the normalized number of communications for tissue-specific metabolite-sensor pairs (x-axis). Tissue labels (y-axis) were aligned with that in **4A**. The warmer the color, the higher the communication number, and vice versa. (D) Violin plots of highly expressed metabolite-sensor pairs in topic 13. The color represented the expression level based on the communication score. A missing violin indicated the metabolite-sensor pair was not expressed in that tissue.

You et al. also found that in ovarian cancer (EOC), SLC7A1 inhibited the expression of the CCL4 that was related to the immune infiltration microenvironment, suggesting a great candidate for EOC therapy [34]. Together, we identified highly tissue-specific CCC topics through topic modeling. The identified CCC events pose the potential to regulate EC heterogeneity across tissue types.

### Topic modeling revealed tissue-specific protein-mediated CCC patterns

Numerous studies have shown that protein ligand-receptor interactions are crucial in regulating cell function and cell fate [35,36]. By applying topic modeling on protein-mediated CCC landscapes inferred by LIANA+ [28,29], we identified 15 CCC topics involving ECs (**Supplementary Table 2**). Interestingly, 13 of 15 protein-CCC topics were uniquely associated with one tissue type (**Fig. 5A**). For example, topic 13 was strongly enriched with the thymus, while topic 15 was strongly associated with the heart. Visualizing the ligand-receptor pairs that participated in those communication events, we spotted that the ligand-receptor pairs were highly topic-specific (**Fig. 5B**), further suggesting the tissue-specific CCC pattern. By checking the number of communications associated with each pair (**Fig. 5C**), many ligand-receptor pairs indeed showed a much higher number of communications in one tissue compared to other tissues, consistent with the patterns observed above (**Fig. 5B**). These results suggested that the identified ligand-receptor pairs in topics were tissue-specific, which indicated their unique, important roles in regulating tissue-specific EC functions. Here, we have dived deeper into topic 15, which was a topic highly related to the heart (**Fig. 5A**). We focused on six ligand-receptor pairs based on their significantly high probability score (topic-related) and high communication number (tissue-related) (**Fig. 5B-C**), respectively. Those ligand-receptor pairs were ADAM17∼ITGB1, CALM1∼RYR2, ADAM17∼ITGA5, LAMA2∼RPSA, ADAM17∼NOTCH1, and MAML2∼NOTCH1. Interestingly, those pairs’ ligand and receptor gene expression were also highly expressed in the heart compared to other tissues (**Fig. 5D**). These identified ligands and receptors have been reported to have important functions in the heart with in vivo and in vitro experiments. For example, ADAM17 was essential to normal cardiovascular functions [37].

**Figure 5:**
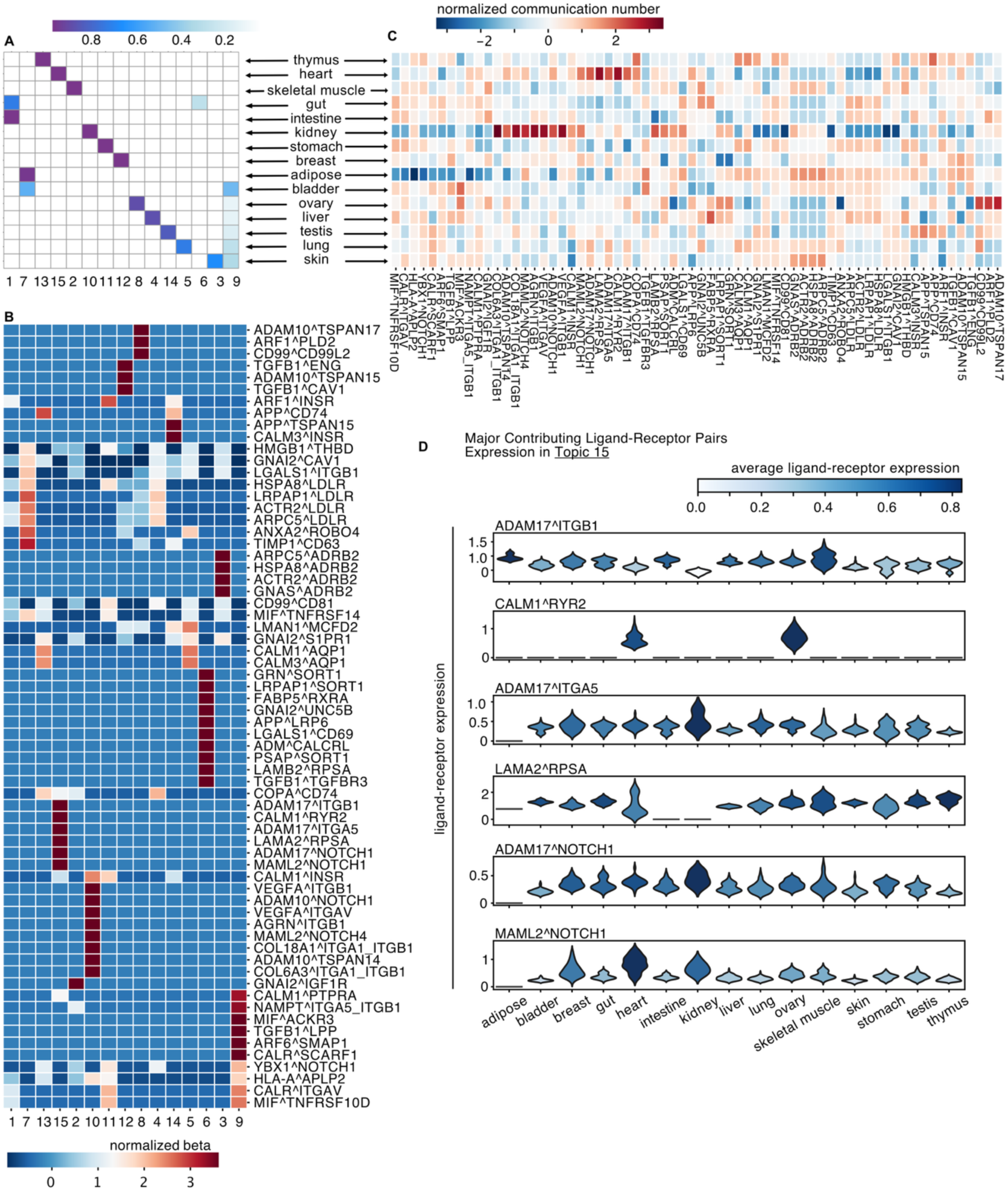
LDA inferred topics of ligand-receptor CCC networks. (A) Clustered heatmap showing the correlation of the LDA results using LIANA+ inferred CCC results. The X-axis indicated inferred topics and the y-axis labeled tissue type. The darker the color, the stronger the correlation between the corresponding topic and tissue. (B) Clustered heatmap of the normalized beta score of selected ligand-receptor pairs (y-axis) in each topic (x-axis). The x-axis label was aligned with that in **5A**. The warmer the color, the higher the specificity, and vice versa. (C) Clustered heatmap of the normalized number of communications for tissue-specific ligand-receptor pairs (x-axis). Tissue labels (y-axis) were aligned with that in **5A**. The warmer the color, the higher the communication number, and vice versa. (D) Violin plots of highly expressed ligand-receptor pairs in topic 15. The color represented the average of the pair expression level. A missing violin indicated the ligand-receptor pair was not expressed in that tissue.

Elevated ADAM17 levels have been found in the lipid core of human atherosclerotic plaques and mouse models with atherosclerotic lesions progression [38]. ADAM17 also plays a crucial role in cardiac inflammation by cleaving and activating various proinflammatory cytokines and their receptors [39]. It shows that ADAM17 is an important ligand that regulates normal heart functions. Moreover, NOTCH1, which interacts with ADAM17 and MAML2 in the heart, was also a crucial receptor in tissue formation and homeostasis. Ye et al. have revealed that the NOTCH1 deficiency could cause defects in the proliferation of early human cardiomyocytes [40]. Roifman et al. found reduced NOTCH1 levels in both ventricular outflow tracts, suggesting that NOTCH1 haploinsufficiency may cause bicuspid aortic valve and related cardiac anomalies [41]. A recent clinical study also showed an association between the NOTCH1 variants and cardiac and extracardiac vascular anomalies [42]. This analysis provided insights into tissue-specific communication patterns to EC for the ligand and receptor pairs and further poses the potential to explain how EC heterogeneity was regulated at the tissue niche level.

## Discussion

Tissue-specific EC functions have been widely studied within individual tissues and compared between vessel types. It is unclear how human EC heterogeneity across different tissue types was regulated systematically. An intriguing question is whether ECs should continue to be grouped by vessel types or should they be further grouped by tissue types.

In this study, we investigated the microenvironments led by CCC as a potential factor to regulate EC differences across vessel and tissue types. Analyzing human EC scRNA-seq data specifically, we first identified different patterns of EC heterogeneity, which were shown by tissue type and vessel type separately. Aside from the well-known knowledge that the EC vessel types differed due to their specific locations and functions, ECs in some tissues, e.g., the heart and kidney, were more different under the tissue context than vessel types. This finding was comparable with other studies. Chen et al. showed that the embryonic EC heterogeneity came from tissue origin and vascular-bed position [43]. They also found that embryonic ECs in some tissues, such as the heart and liver, expressed tissue-selective markers.

In contrast, ECs in other tissues were more similar within each vessel type. Other groups also identified EC heterogeneity, as shown by tissue-specific marker genes and enriched functions, but with the mouse model [9,10]. In this study, we dived deeper into the microenvironment regulation by CCC and found tissue-specific CCC patterns for EC in humans. We investigated EC communications under various tissue contexts using an unsupervised machine-learning algorithm and topic modeling. For metabolite-mediated and protein-mediated communications, many CCC events inferred *in silico* were exclusively strong in one tissue compared to other tissues. Based on the previous reports, we further verified that the identified ligands and receptors were associated with essential functions in specific tissues. In summary, the identified CCC events in each topic were promising candidates for further investigation to provide insights into EC tissue-specific differences regulated by metabolite-mediated and protein-mediated CCC.

## Method and materials

### Data acquisition and process

We downloaded over 3,000,000 single cells of 15 human tissues from the DISCO database[44], including adipose, bladder, breast, gut, heart, intestine, kidney, liver, lung, ovary, skeletal muscle, skin, stomach, testis, and thymus. The downloaded data in each tissue includes raw gene expression counts and cell metadata annotation, such as cell type, project identification number, disease status, etc. First, we filtered out disease samples to focus on data from normal tissues, which resulted in over 2,000,000 single cells of 15 tissue types. The pre-processing, including normalization, was done in Python by Scanpy (version 1.8.2) based on the raw count matrix. The normalized data was used for the downstream analysis in this study, including EC heterogeneity analysis and cell-cell communication analysis. In the EC heterogeneity analysis, we retrieved single cells with cell types labeled as four major EC vessel types, namely, arterial, capillary, lymphatic, and venous EC. We finally removed the batch effect using the Harmony algorithm wrapped in the Scanpy package.

### Dimensionality reduction and clustering

We did uniform manifold approximation and projection (UMAP), reducing the dimensionality to show the clustering patterns across EC vessels and tissue types. We further investigated the difference between EC vessel type versus tissue type by performing principal component (PC) analysis using the Harmony-corrected PCs (50 in total). Results were visualized using PC1 and PC2, which explained the most variance. Finally, hierarchical clustering based on tissue type and vessel type was done with Scipy using 50PCs; the ward variance minimization algorithm and Euclidean metric were used to build the model. Pairwise distance (Euclidean) was calculated using the scikit-learn package for two major clusters for further quantifications.

### Cell-cell communication analysis

To increase the computation efficiency, we down-sampled data from each tissue to contain only 10,000 cells and did normalization. Then, we utilized two bioinformatics tools to infer cell-cell communications. MEBOCOST [27] predicted metabolite-mediated CCC events. LIANA+ [28,29] was used to predict protein-mediated CCC. Each inference was performed on cell types within each tissue type. Default parameters were used when running the framework. The input data contained all cell types. We focused on communication events with a false discovery rate (FDR) of less than 0.05. Then, we restricted the predicted communications with the type of sender/source or receiver/target as ECs to further focus on EC-related CCC for the downstream analysis.

### Topic modeling

Focusing on how EC was regulated, we further filtered our data to only allow EC as the receiver/target cells in a single communication event. Then, topic modeling was performed with Latent Dirichlet Allocation (LDA) in R. The input matrix was tissue type by communication events, filling with the number of communication events predicted in each tissue. The number of topics in the LDA analysis was set at 15 based on the assumption that each tissue formed its topic. The top 10 communication pairs (with the highest beta score from topic modeling) were selected to visualize topic-specific patterns due to their strong probability in each topic. The topic’s word distribution, beta β, and communication number for each metabolite-sensor/protein-receptor pair were normalized for each topic and tissue, respectively. To validate the tissue-specific communication events, the communication score (product of the average metabolite and sensor expression) from the MEBOCOST and the average expression of ligand-receptor pairs from the CellPhoneDB were used, respectively.

### Quantification and statistical analysis

Statistical analyses were performed using Python 3.10. The Euclidean distance matrix was used to calculate the correlation. The data were presented as mean in bar plots, and the box plots displayed quantiles and median.

### Code and data availability

All the analyses were done using Python (version 3.8 for raw data processing and 3.10 for downstream analysis), except for the LDA analysis, which was performed using R (version 4.4.0). The script to reproduce all analyses in this study can be found at https://github.com/zhuzimoo/EC_project.

## Acknowledgments

During the period of this project, K. C. is supported in part by the grants R01GM125632, R01GM138407, R01HL148338, R01HL133254, R01CA278832, and R01HL174928 to K. C., L. Z. is supported in part by the grants R01HL155632 to L. Z. and Additional Ventures (Single Ventricle Research Foundation) to L. Z, R. Z is supported in part by the grants K99HG013662 to R. Z. and NextGen Accelerator Award (Cell Discovery Network).

## Author Contributions

K. C. and R. Z. conceived the project. Z. Z. performed all the analyses in this study and drafted the manuscript. All other co-authors read and revised the manuscript.

