## Supplementary Fig. for "Human single-cell atlas analysis reveals heterogeneous endothelial signaling"

### Supplementary Figure 1

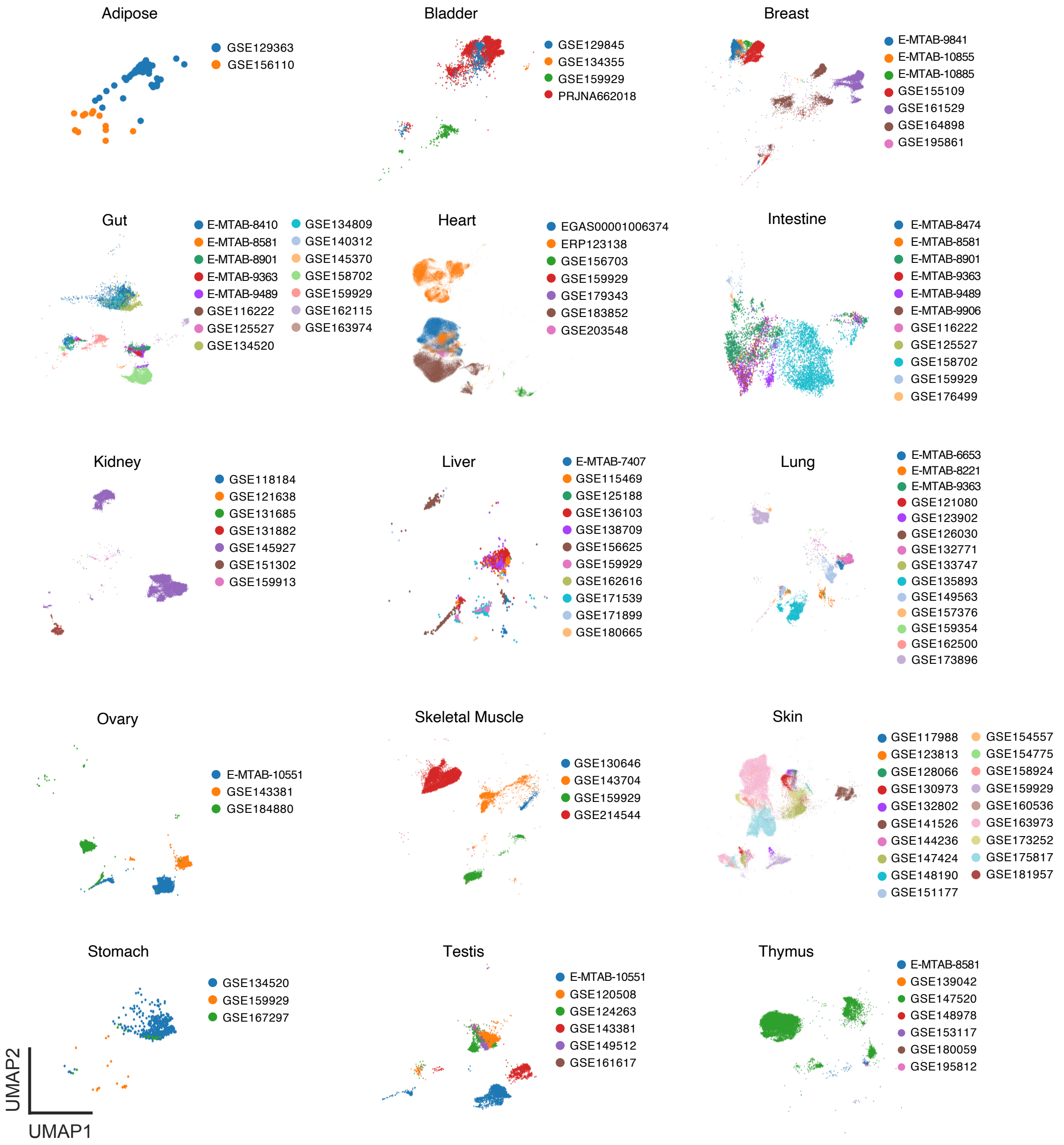


**Supplementary Figure 1: Batch effect of UMAP clustering with raw scRNA-seq data**

UMAP projections with color-coded different projects from 15 tissues.

### Supplementary Figure 2

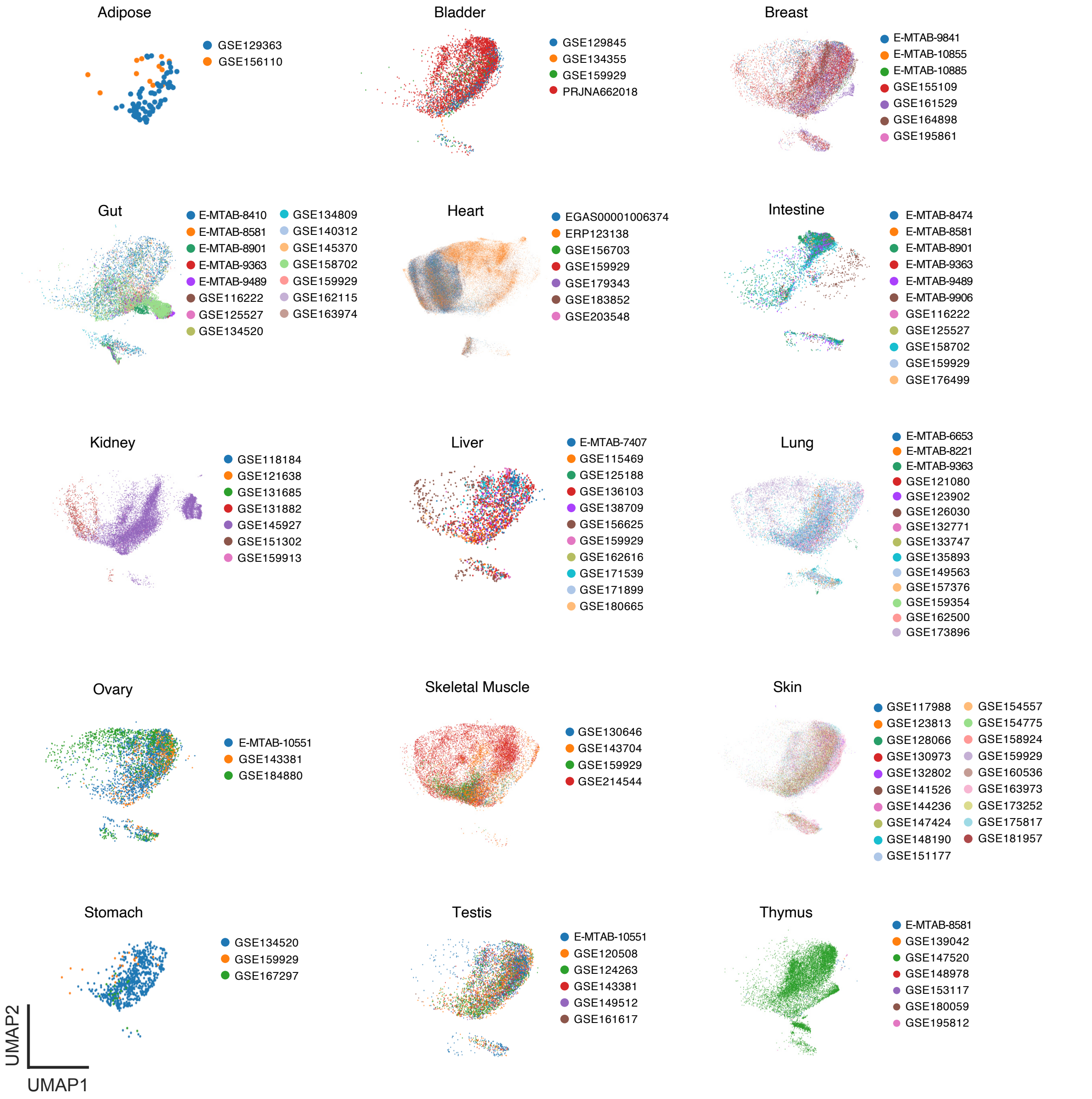


**Supplementary Figure 2: UMAP clustering after batch effect removal**

UMAP projections with color-coded different projects from 15 tissues. The colors used for each tissue were consistent with that of Supplementary Fig.2.

**Supplementary Table 1: The metadata of scRNA-seq data used in this study.** Tissue name, project accession, and number of cells were listed in the table.

**Supplementary Table 2: Top ranked ligand-receptor and metabolite-sensor pairs in each CCC topic.** The spreadsheet of "met-sensor" includes top presented metabolite-sensor pairs in each CCC topic along with the probability score (beta value). The spreadsheet of "protein-ligand" includes top presented ligand-receptor pairs in each CCC topic along with the probability score (beta value)
